# Mutational analysis to explore long-range allosteric coupling and decoupling in a pentameric channel receptor

**DOI:** 10.1101/2020.07.03.173997

**Authors:** Solène N. Lefebvre, Antoine Taly, Anaïs Menny, Karima Medjebeur, Pierre-Jean Corringer

## Abstract

Pentameric ligand-gated ion channels (pLGICs) mediate chemical signaling through a succession of allosteric transitions that are yet not completely understood. On the prototypic bacterial channel GLIC, we explored the conformational landscape of the protein during pH-gating. To this aim, we introduced a series of allosteric mutations, and characterized the protein conformation over a broad pH range. We combined electrophysiological recordings, fluorescence quenching experiments monitoring key quaternary reorganizations, and simulations by normal mode analysis. Moderate loss-of-function mutations and the allosteric modulator propofol displace allosteric equilibria involved in pre-activation and pore opening processes, highlighting long-range allosteric coupling between distant regions of the protein. In contrast, total loss-of-function mutations stabilize the protein in unique intermediate conformations where motions are decoupled. Altogether, our data show that the protein can access a wide conformational landscape, raising the possibility of multiple conformational pathways during gating.

## Introduction

Pentameric ligand gated ion channels (pLGICs) mediate fast synaptic communication in the brain. In mammals, this family includes the excitatory nicotinic acetylcholine and serotonin receptors (nAChRs and 5-HT_3_Rs) as well as the inhibitory γ-aminobutyric acid (GABA) and glycine receptors (GABA_A_Rs and GlyRs) (Jaiteh et al., 2016). pLGICs are also present in bacteria, notably with the pH-gated channels GLIC (Bocquet et al., 2007) and sTeLIC (Hu, Nemecz, et al., 2018), the GABA-gated channel ELIC (Zimmermann and Dutzler, 2011), and the calcium-modulated DeCLIC (Hu et al., 2020).

pLGICs physiological function is mediated by alternating between different allosteric conformations in response to neurotransmitter binding. Seminal work in the 80s showed that a minimal four-state model describes the main allosteric properties of the muscle-type nAChR (Heidmann and Changeux, 1980; Sakmann et al., 1980). The ability of ACh binding to activate the nAChR involves a resting-to active-state transition. In addition, prolonged ACh occupancy promotes a biphasic desensitization process underlying the transition to so-called fast- and slow-desensitized states. Subsequently, kinetic analysis of the close-to-open transitions recorded by single channel electrophysiology unraveled multiple additional states that are required to account for the observed kinetic patterns. For activation, short-lived intermediate “pre-active” states named “flipped” (Lape et al., 2008) and “primed” (Mukhtasimova et al., 2009) were included in the kinetic schemes of the GlyRs and nAChRs, while rate-equilibrium free-energy relationship (REFER) analysis of numerous mutants of the nAChR suggested passage through four brief intermediate states (Gupta et al., 2017). Likewise, analysis of single-channel shut intervals during desensitization are described by the sum of four or five exponential components, suggesting again additional intermediate states (Elenes and Auerbach, 2002). Kinetics data thus show that pLGICs undergo a complex cascade of structural reorganizations in the course of both activation and desensitization. Those events are at the heart of the protein’s function, allowing the coupling between the neurotransmitter site and the ion channel gate which are separated by a distance of 5 nm in pLGIC structures.

The past decade has seen great advances by structural biology to seek understanding about the molecular mechanisms involved in gating (Nemecz et al., 2016). At least one structure of each major member of prokaryotic (Hilf and Dutzler, 2008; Bocquet et al., 2009; Hu, Nemecz, et al., 2018; Hu et al., 2020) and eukaryotic pLGICs (Althoff et al., 2014; Du et al., 2015; Polovinkin et al., 2018; Gharpure et al., 2019; Masiulis et al., 2019) have been resolved by X-ray crystallography or cryo-EM. They highlight a highly conserved 3D architecture within the family. Each subunit contains a large extracellular domain (ECD) folded in a β-sandwich and a transmembrane domain (TMD) containing four α-helices, with the second M2-helix lining the pore. However, the physiological relevance of structures or their assignment to particular intermediates or end-states in putative gating pathways remains ambiguous and poorly studied. Conversely, it is likely that key conformations, unfavored by crystal packing lattice or under-represented in receptor populations on cryo-EM grids, are missing in the current structural galleries.

Understanding the allosteric transitions underlying gating thus requires complementary techniques, where the protein conformation can be followed in near-physiological conditions, i.e. at non-cryogenic temperature on freely moving protein, and over a broad range of ligand concentrations. To this aim, we previously developed the Tryptophan/Tyrosine induced quenching technique (TrIQ) (Menny et al., 2017) on GLIC, a proton-gated channel (Parikh et al., 2011; Laha et al., 2013; Gonzalez-Gutierrez et al., 2017). In this technique, the protein is labeled with a small fluorophore, the bimane, and collisional quenching by a neighboring indole (tryptophans) or phenol (tyrosines) moieties is used to report on changes in distance between two residues within the protein (Mansoor et al., 2002, 2010; Jones Brunette and Farrens, 2014). Bimane/quencher pairs on GLIC combined with kinetic analysis allowed us to characterize pre-activation motions occurring early in the conformational pathway of activation (Menny et al., 2017). Indeed, they occur at lower proton concentrations than pore opening, and are complete in less than a millisecond, much faster that pore opening that occurs in the 30-150 millisecond range in electrophysiology (Laha et al., 2013).

Here, we exploit the TrIQ approach to explore the conformational landscape of GLIC during pH-gating, in combination with allosteric ligands and mutations. To help interpreting the fluorescence quenching data into structural terms, we first built atomistic models of the various bimane-labeled proteins, and simulated their gating transition pathways using coarse grain modeling and normal mode analysis. We then performed electrophysiology and fluorescence quenching experiments on a series of GLIC mutants presenting impaired activity. Our results indicate that mutations alter the function by distinct mechanisms, either displacing the allosteric equilibriums already present in the wild-type protein, or stabilizing the protein into novel non-conducting states, suggesting multiple conformational pathways during gating.

## Results

### Normal Modes Analysis generates two distinct conformational pathways for GLIC activation

We first performed atomistic simulations to characterize the protein motions associated with the transition between resting and active conformations. To do so we modeled the transition path between GLIC-pH7 (4NPQ) and GLIC-pH4 (4HFI) X-ray structures using iMODfit (Lopéz-Blanco and Chacón, 2013). This program based on Normal Mode Analysis (NMA) originally designed to perform flexible fitting of atomic coordinates in cryo-EM densities, has also been successfully used to investigate concerted motions on NMDA receptors (Esmenjaud et al., 2019). GLIC-pH7 is in a non-conductive conformation with a closed hydrophobic gate in the upper part of the pore, consistent with a resting-like state. The GLIC-pH4 structure shows in contrast an open gate compatible with a conductive conformation (Cheng et al., 2010; Fritsch et al., 2011; Sauguet et al., 2013; Gonzalez-Gutierrez et al., 2017), consistent with an active-like structure.

Two trajectories, A and B, were computed between the two end state structures and divided into 12 and 11 frames respectively. RMSD analysis between each frame and the target structure underlie gradual reorganization of GLIC across the length of both simulations (Figure 1A). Both trajectories, when visualized from the resting to active state, show three major components of re-organizations, namely a quaternary twist of the pentamer, a reorganization of the pore region leading to its opening/closure, and a quaternary compaction of the ECD.

**Figure 1.**
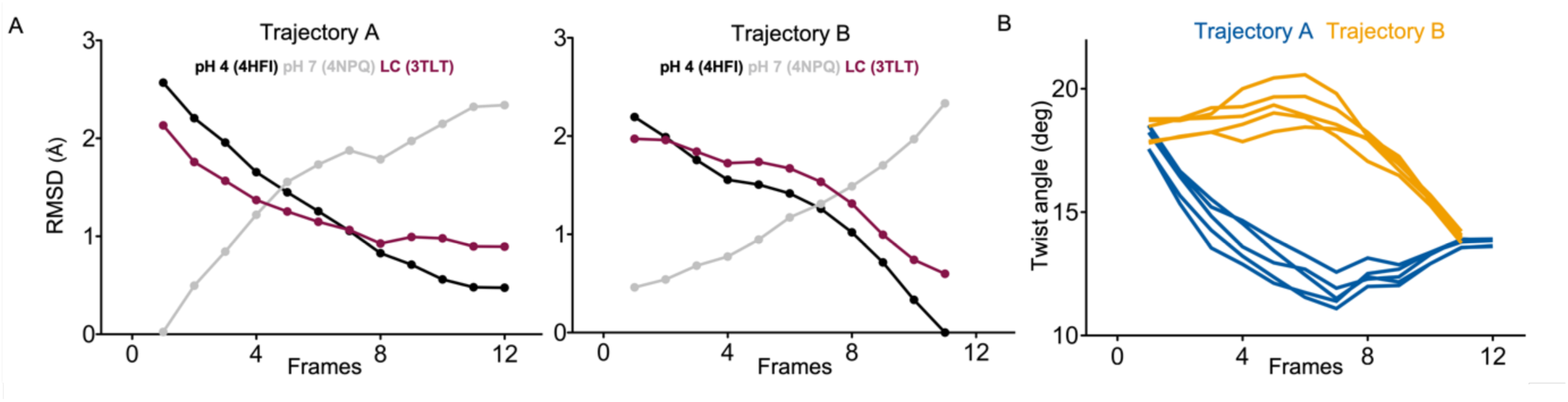
iMODfit generates two distinct trajectories for GLIC activation. (A) RMSD evolution throughout the frames of the A and B trajectories against GLIC structures at pH4 (black); pH7 (gray) and locally-closed (LC; purple), pdb codes are indicated. Both trajectories are shown with frame 1 being the closest to GLIC-pH7. (B) Twist angle measured throughout the frames on both trajectories. The twist angle is measured by the angle formed between vectors from the center of mass of the protein and the centers of mass of the ECD and TMD as defined in (Calimet et al., 2013). Each trace corresponds to the trajectory of a single subunit within the pentamer.

In the A trajectory, the twist motion occurs in the first half of the trajectory (Figure 1B). This motion describes opposite rotations between ECD and TMD domains, as measured by the twist angle defined by center of mass vectors of the ECD and TMD (Taly et al., 2005; Calimet et al., 2013). The reorganization of the pore happens in the second half of the trajectory and leads to the opening of the hydrophobic gate, as evaluated by the radius at the level of Ile233 (also named I9’; Figure 2A & 2B). This motion is associated with a tilt of M2 toward M3 (as measured by a decrease in distance between His235 nitrogen and the carbonyl backbone of Ile259 that are at H-bond distances in the GLIC-pH4 structure; Figure 2A & 2C), and with the outward motion of the M2-M3 loop (monitored by an increase in distance between the Cα of residue Pro250 and phenolic oxygen of Tyr197; Figure 2A & 2D). Pore opening is also associated with, at the bottom of the ECD, a contraction of the β-sandwich (evaluated by a decrease in distance between C_α_ of residues Asp32 and Gly159; Figure 3A & 3C), as well as quaternary reorganization around loop 2 (measured by a decrease in inter-subunit distance C_β_ Lys33/Trp160, Figure 3-Supplement 1). In addition to these two consecutive global motions, the progressive quaternary compaction of the ECD, another crucial landmark of GLIC reorganization, occurs throughout the trajectory. This compaction is quantified through measurement of inter-subunit distances (between C_β_ Asp136/Gln101 and Arg133/Leu103; Figure 3A, 3B & 3-Supplement 2), indicating a progressive decrease in distance throughout the frames. It is noteworthy that these inter-subunit distances are highly variable, due to the asymmetric nature of the ECDs of the GLIC-pH7 structure, where each subunit β-sandwich presents a unique orientation as well as relatively high B-factors (Sauguet et al., 2014). This variability decreases over the frames to reach the structure of GLIC-pH4 which is compact and essentially symmetric.

**Figure 2.**
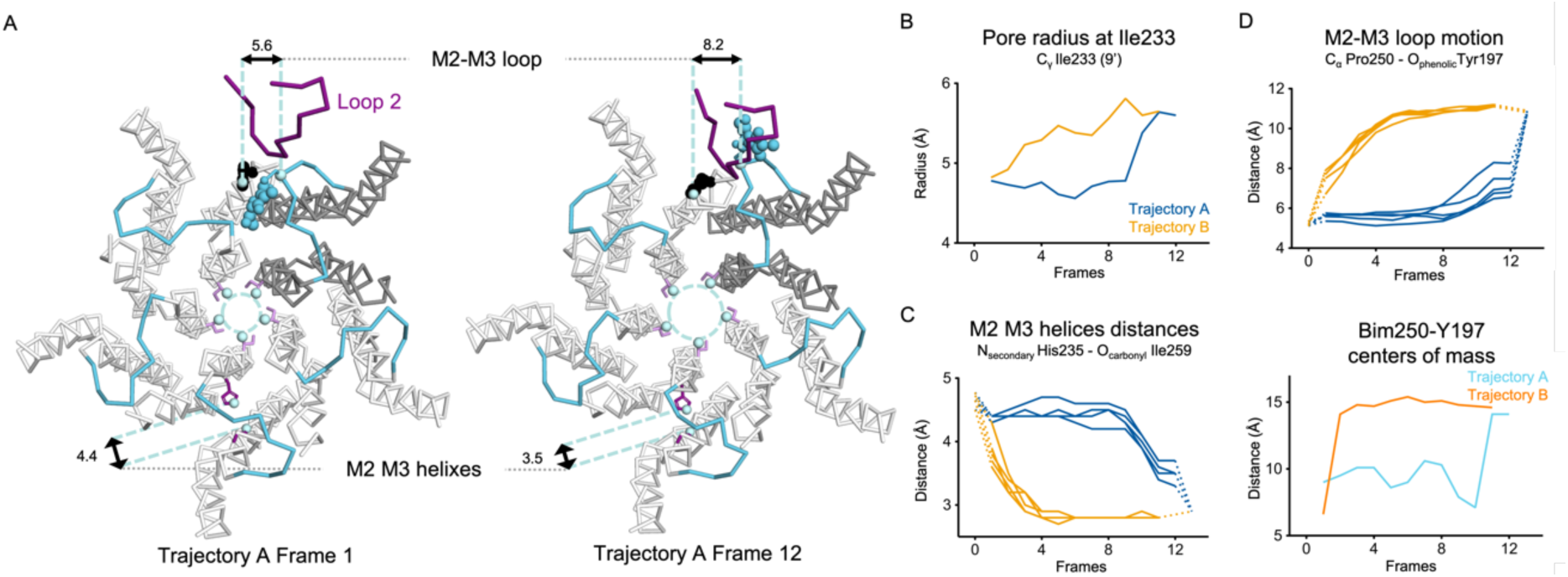
Key motions of the TMD in A and B trajectories. (A) Snapshots of GLIC TMD top view in the first and last frame of the A trajectory with a Bim250-Y197 quenching pair modeled. Bimane is shown in blue and quencher in black spheres. One subunit is shown in grey, the other are in white, the M2-M3 loop is shown in blue and the loop 2 from ECD is shown in purple for one subunit. Atoms used for measurements are shown in pale blue spheres and distances are indicated in angstroms. (B) Pore radius measured at the Ile233 level. (C) Intra-subunit separation of M2 and M3 helices measured between atoms indicated. Points at frames 0 and 13 are the distances measured in pH4 and pH7 X-ray structures. (D) Inter-subunit distances showing M2-M3 loop outward motion at the Pro250-Tyr197 level (top panel) and between bimane and Tyr197 centroids (bottom panel) in both trajectories A and B.

**Figure 3.**
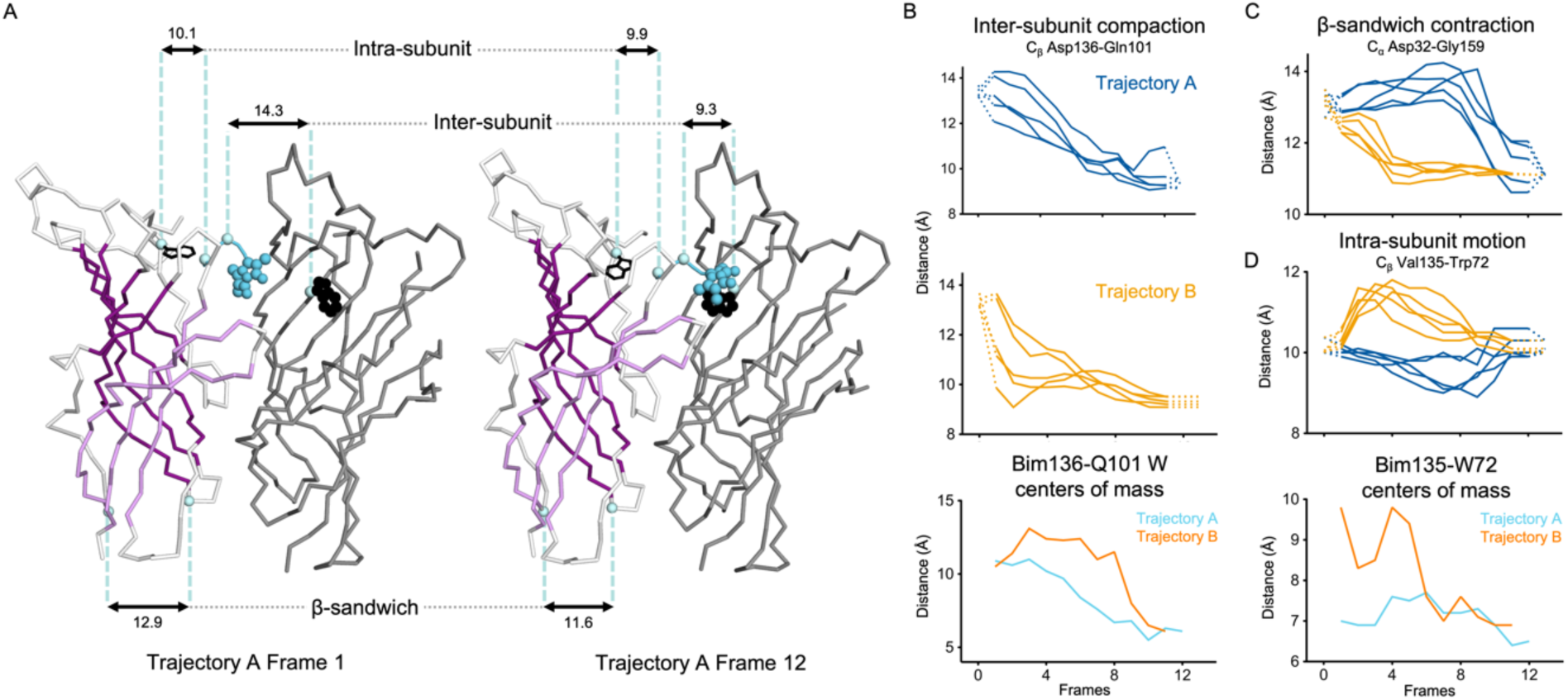
Key motions of the ECD in A and B trajectories. (A) Snapshots of two subunits of GLIC ECD in the first and last frame of the A trajectory with a Bim136-Q101W quenching pair modeled. One subunit is shown in grey, the other in white with sheets of the β-sandwich shown in dark and light purple; bimane is shown in blue and quencher in black spheres; C_α_ and C_β_ atoms used for measurements are shown in pale blue spheres and distances are indicated in angstroms. (B) Inter-subunit distances showing ECD compaction at the Asp136-Gln101 level (first two panels) and between bimane and Q101W centroids (bottom panel) in both trajectories A and B. Points at frames 0 and 13 are the distances in pH4 and pH7 X-ray structures. (C) Intra-subunit distance showing contraction at the bottom of the β-sandwich measured by C_α_ distances between Asp32 and Gly159. (D) Intra-subunit C_β_ distances between Val135 and Trp72 (top panel) and between Bim135 and Trp72 centers of mass (bottom panel).

Trajectory B shows substantially the same components but with an inverted sequence of events, pore opening and associated motions start first, followed by the twist motion in the last frames, the compaction of the ECD being spread over the whole trajectory. In conclusion, iMODfit generate two distinct trajectories that are equally plausible to describe a gating transition of activation of GLIC.

### Combined quenching-docking data establish Bim136-Q101W and Bim250-Y197 as reporter of key allosteric motions of GLIC

To directly relate the conformational reorganizations generated by iMODfit to the previously collected fluorescence quenching data (Menny et al., 2017), we modeled the various fluorophore/quenching pairs in both trajectories. In the fluorescence experiments, a bimane fluorophore was covalently labelled on a cysteine at the indicated position (after mutation of the endogenous cysteine C27S), and a Trp or Tyr quenching residue was incorporated when necessary. We focused our analysis on Bim136-Q101W, Bim135-W72 and Bim250-Y197 (Bim133-L103W and Bim33-W160, not used in following fluorescence experiments, are presented in Figure 3-Supplementary 1 & 2). For modeling of each pair, the cysteine mutations (and quencher when necessary) were performed and the bimane moiety was docked into each frame while keeping its carbon at a covalent-bond compatible distance to the sulfur atom of the cysteine. The distance between the centers of mass of the bimane and indole/phenol moieties in each frame was then measured to follow the evolution of the distance between the bimane and quencher.

For the ECD quenching pair Bim136-Q101W, the simulations show that Bim136 and the indole ring of Trp101 are separated in the resting-like state, and are in close contact in the active-like state (Figure 3A). These observations are in good agreement with the previously published fluorescence data that show a decrease in fluorescence intensity upon pH drop indicating a decrease in distance between the bimane and quencher. The A trajectory shows a progressive decrease in distance that parallels the ECD quaternary compaction movement described previously (C_β_ Asp136/Gln101), whereas the B trajectory shows a more variable pattern, and a sharper distance decrease only in the last frames (Figure 3B).

For the ECD-TMD interface quenching pair Bim250-Y197, our simulations show that Bim250 is in close contact with the phenol ring of Tyr197 in the resting-like state, while both moieties are separated in the active-like state, the bimane moiety moving on the other side of loop 2 (Figure 2A). This is also in agreement with the fluorescence recordings that showed an increase in fluorescence upon pH-drop indicating that the Bim250 is moving away from its quencher Tyr197. Both A and B trajectories show an abrupt change in Bim250/Y197 distances, corresponding respectively to a late versus early separation, and these changes occur during the outward motion of the M2-M3 loop (P250_Cα_/Y197_O_ distance; Figure 2D).

In contrast, for Bim135-W72 (Figure 3D), the distances between the centers of mass of the bimane and indole moieties are not correlated with the distances between the residue’s backbone, while the pH-dependent changes in fluorescence show a bell shape curve suggesting important but complex changes in distances at this level. Since at this position bimane occupies a rather buried location within the protein structure, we suggest that these discrepancies come from local reorganization of surrounding residues that are not directly correlated with the movement of the backbone. Interestingly, we previously solved the X-ray structure of Bim135-W72 at pH4 by crystallography, which shows a similar location of the bimane moiety with that of our docking in GLIC-pH4 (Figure 3-Supplementary 3).

In conclusion, for Bim136-Q101W and Bim250-Y197 quenching pairs, the simulations of the end states show good agreement with steady-state fluorescence data. Interestingly, kinetic analysis show that the conformational motions monitored by both pairs report on a pre-activation transition that occurs well before channel opening (Figure 4A; Menny et al., 2017). This observation is compatible with trajectory A, for which the quenching of Bim136-Q101W occurs throughout the trajectory from frame 1 to 10, the unquenching of Bim250-Y197 occurs between frame 10 and 11, while the full reorganization of the pore and associated motions are completed only in the very last frame of the trajectory. The pre-activation scheme is clearly not compatible with trajectory B, that shows an inverted order of events.

**Figure 4.**
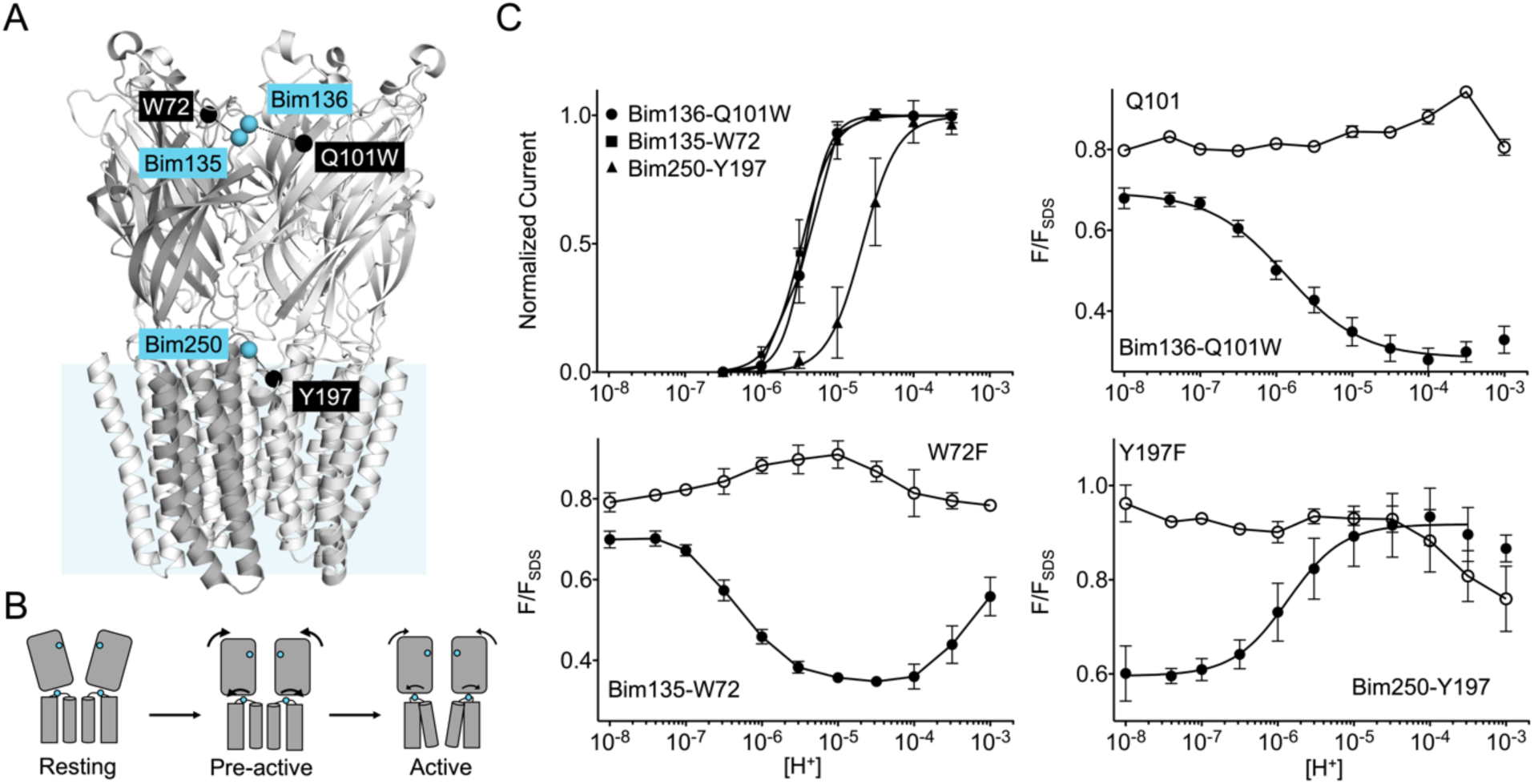
Quenching pairs used in the study. (A) GLIC-pH 4 structure side view. Blue spheres show the Cα of the residues that were submitted to cysteine mutation and bimane labeling (Bim135, Bim136 and Bim250), and black spheres Cα of the quenchers (W72, Q101W and Y197). (B) Pre-activation scheme for GLIC activation, showing a first pre-activation step involving full compaction of the ECD and motion of the M2-M3 loop as monitored by fluorescence, followed by a pore opening step. (C) pH-dependent response curves of the three mutants labelled with bimane by electrophysiology and bimane fluorescence. Fluorescence data are shown normalized to the fluorescence of the denatured protein (F_SDS_), bimane fluorescence is shown without quencher (empty points) and in presence of the quencher (filled points).

To explore the conformational landscape of GLIC and of its mutants, we thus monitored in parallel the ECD quaternary compaction with the Bim136-Q101W fluorescent sensor and reorganizations at the TMD level using electrophysiology or the fluorescent sensor Bim250-Y197 reporting on M2-M3 loop motions. In specific cases, we additionally used Bim135-W72 to monitor tertiary motions in the ECD.

### The quaternary compaction of the top of the ECD is strongly allosterically coupled with the lower part of the ECD interface

To allow an accurate comparison between mutants, we first measured detailed pH-dependent fluorescence and electrophysiological curves of the three selected bimane-labelled GLIC (i.e. Bim136-Q101W; Bim250-Y197 and Bim135-W72, Figure 4B). Fluorescence was measured in steady-state conditions on detergent (DDM)-purified protein, and normalized to the fluorescence intensity under denaturing conditions (1% SDS), as previously described (Menny et al., 2017). GLIC allosteric transitions are indeed particularly robust in different lipid/detergent conditions (Sauguet et al., 2014; Carswell et al., 2015) and DDM-purified protein yielded similar results to that of asolectin-reconstituted protein (Menny et al., 2017), while allowing better reproducibility. For the three pairs, we confirmed that the pH-dependent fluorescence changes are essentially abolished when mutating the quenching partner (Figure 4C), phenylalanine not being able to quench bimane fluorescence (Mansoor et al., 2002). We actually identify here Trp72 as the endogenous quenching residue of Bim135 since the mutant Bim135-W72F is functional in electrophysiology, but does not undergo pH-dependent quenching (figure 4C). We also confirm that pH-dependent quenching curves for Bim136-Q101W and Bim250-Y197 display higher sensitivity (especially for Bim250-Y197) and lower cooperativity than the pH-dependent activation curves recorded by electrophysiology (Figure 4C).

We first investigated allosteric mutants located at the inter-subunit interface in the lower part of the ECD (Figure 5A). We previously showed that E26Q produces a decrease in pH_50_ for activation (Nemecz et al., 2017), a phenotype which is conserved here on the Bim136-Q101W background (Figure 5B and 5C). The fluorescence quenching curve of the Bim136-Q101W pair also shows a decrease in pH_50_ in presence of the E26Q mutation (Figure 5D), and the ΔpH_50_ between WT and E26Q are nearly identical in electrophysiology and fluorescence (−0.59 and 0.57 respectively; Table 1). Interestingly, the fluorescence quenching curve of E26Q has a remarkable feature as compared to all other mutants investigated thereafter: in the pH7-8 range, where the pH-dependent fluorescence quenching is not yet observed, the fluorescence (F/F_SDS_) is significantly lower (F_0_ = 0.53) as compared to the Bim136-Q101W (F_0_ = 0.71). This suggests that substantial quenching is present at neutral pH and E26Q not only alters the allosteric transition, but also modifies the conformation of the resting state itself which appears to be more compact when the E26Q mutation is present.

**Table 1.** Dose-dependence of electrophysiological and fluorescence quenching responses. pH_50_ and Hill coefficient n_H_ average and standard deviation values are shown after individual fitting of all measurements. Number of experiments n correspond to the number of oocytes for electrophysiology and the number of fluorescence measurements, each measurement including values for a full pH range. F_0_ corresponds to the initial fluorescence value at pH 7/8 and F_max_ the maximum variation in fluorescence amplitude within the pH range. ΔpH_50_ values are calculated in comparison with mutants and their parent construct Bim136-Q101W or Bim250-Y197. NF stands for non-functional and ND for not determined. Hill coefficients have been constrained to 2.5 or below 3 in order to reasonably fit Bim136 + Propofol current and Bim136 + Y28F fluorescence respectively.

| Mutant | Electrophysiological response<br>bimane labeled |  |  |  | Fluorescence quenching response<br>in detergent solution |  |  |  |  |  |
| --- | --- | --- | --- | --- | --- | --- | --- | --- | --- | --- |
| | $pH_{50}$ | $n_H$ | n | $\Delta pH_{50}$ | $pH_{50}$ | $F_0$ | $F_{max}$ | $n_H$ | n | $\Delta pH_{50}$ |
| Bim136-Q101W C27S | $5.42 \pm 0.08$ | $2.68 \pm 0.33$ | 10 | - | $5.85 \pm 0.21$ | $0.71 \pm 0.03$ | $0.45 \pm 0.06$ | $0.77 \pm 0.18$ | 17 | - |
| +E26Q | $4.83 \pm 0.12$ | $2.98 \pm 0.64$ | 6 | -0.59 | $5.28 \pm 0.34$ | $0.53 \pm 0.02$ | $0.22 \pm 0.02$ | $1.13 \pm 0.27$ | 4 | -0.57 |
| +Y28F | $3.88 \pm 0.08$ | $2.63 \pm 0.68$ | 3 | -1.54 | $3.68 \pm 0.34$ | $0.70 \pm 0.01$ | $0.38 \pm 0.09$ | < 3 | 3 | -2.17 |
| +Y28F & C27 | $5.34 \pm 0.11$ | $2.03 \pm 0.12$ | 6 | -0.08 | ND | ND | ND | ND | - | - |
| +D32E | $4.65 \pm 0.12$ | $2.62 \pm 1.51$ | 6 | -0.77 | $5.52 \pm 0.05$ | $0.68 \pm 0.02$ | $0.38 \pm 0.01$ | $0.75 \pm 0.05$ | 3 | -0.33 |
| +E222Q | $4.68 \pm 0.09$ | $2.51 \pm 0.33$ | 5 | -0.74 | $5.36 \pm 0.14$ | $0.66 \pm 0.03$ | $0.40 \pm 0.04$ | $0.74 \pm 0.18$ | 3 | -0.49 |
| +H235Q | $4.04 \pm 0.21$ | $1.19 \pm 0.31$ | 6 | -1.38 | $5.00 \pm 0.09$ | $0.70 \pm 0.01$ | $0.42 \pm 0.01$ | $0.78 \pm 0.10$ | 3 | -0.85 |
| +Propofol | $5.16 \pm 0.13$ | = 2.5 | 3 | -0.26 | $5.33 \pm 0.06$ | $0.67 \pm 0.02$ | $0.38 \pm 0.01$ | $1.21 \pm 0.21$ | 4 | -0.52 |
| +H235Q & propofol | $4.71 \pm 0.15$ | $1.53 \pm 0.48$ | 6 | -0.71 | $5.67 \pm 0.14$ | $0.70 \pm 0.01$ | $0.43 \pm 0.01$ | $1.25 \pm 0.04$ | 3 | -0.18 |
| +H235F | NF | NF | 3 | - | $5.25 \pm 0.08$ | $0.70 \pm 0.05$ | $0.38 \pm 0.04$ | $1.06 \pm 0.17$ | 3 | -0.60 |
| +L157A | NF | NF | 3 | - | $5.42 \pm 0.65$ | $0.67 \pm 0.01$ | $0.19 \pm 0.09$ | $0.71 \pm 0.46$ | 4 | -0.43 |
| +L246A | NF | NF | 3 | - | $4.87 \pm 0.14$ | $0.65 \pm 0.01$ | $0.24 \pm 0.01$ | $0.79 \pm 0.12$ | 3 | -0.98 |
| Bim250-Y197 | $4.66 \pm 0.18$ | $2.20 \pm 0.54$ | 12 | - | $5.83 \pm 0.17$ | $0.59 \pm 0.04$ | $0.33 \pm 0.09$ | $1.19 \pm 0.28$ | 8 | - |
| +H235F | NF | NF | 3 | - | $5.40 \pm 0.13$ | $0.54 \pm 0.01$ | $0.20 \pm 0.01$ | $1.16 \pm 0.15$ | 4 | -0.43 |
| +L157A | NF | NF | 3 | - | $5.81 \pm 0.19$ | $0.49 \pm 0.06$ | $0.45 \pm 0.07$ | $0.64 \pm 0.19$ | 3 | -0.02 |
| +L246A | NF | NF | 3 | - | $5.53 \pm 0.03$ | $0.64 \pm 0.01$ | $0.30 \pm 0.01$ | $1.69 \pm 0.29$ | 3 | -0.3 |

**Figure 5.**
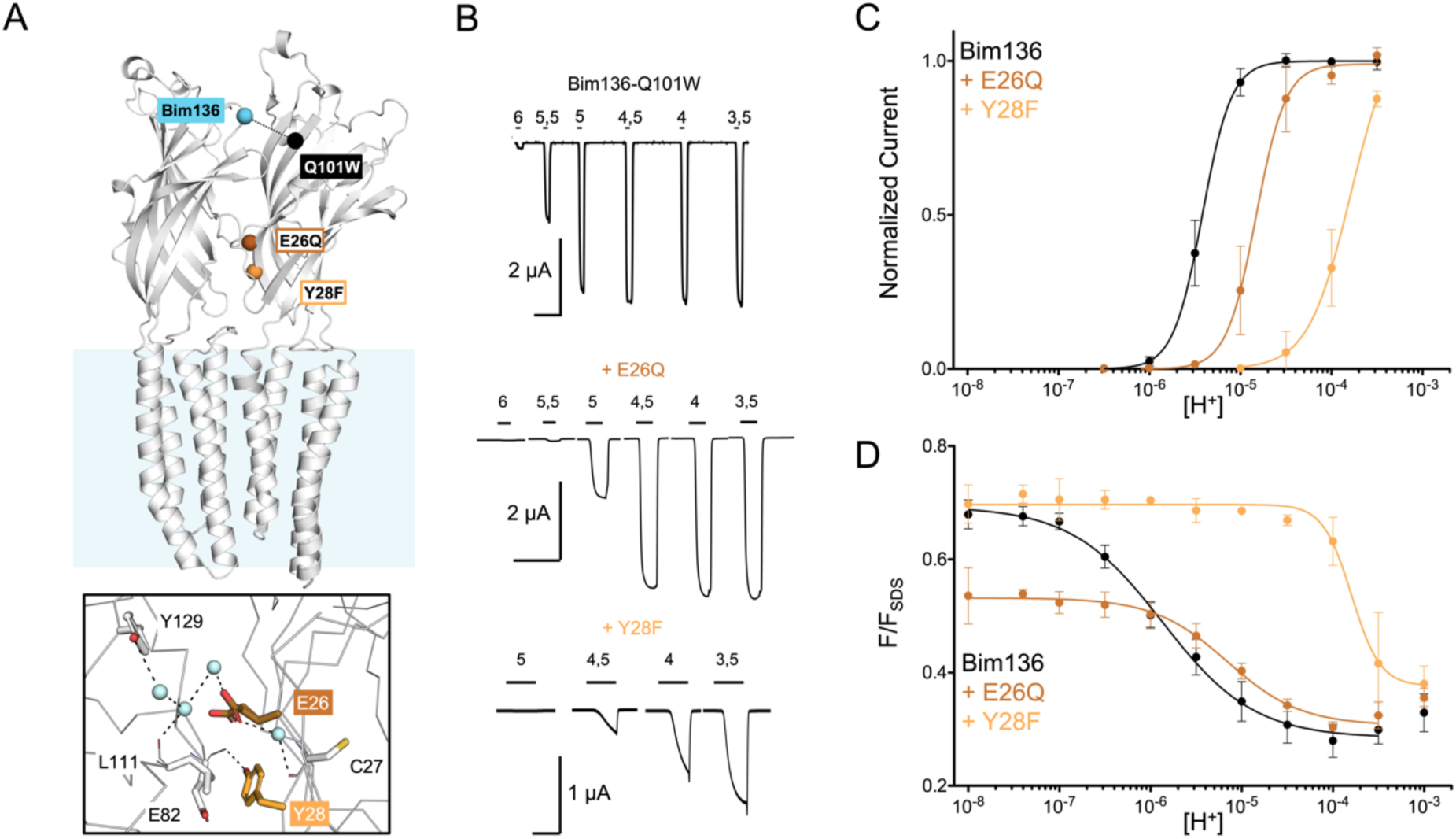
Allosteric coupling within the ECD. (A) Structure of two monomers of GLIC pH 4 showing positions of the fluorescence sensor (Bim136-Q101W) and the two mutations at the bottom of the ECD resulting in a partial loss of function. Lower panel shows a zoom on the interface with Glu26 and Tyr28 residues and their interactions with surrounding residues and water molecules. (B) Electrophysiological recordings in oocytes of the mutants labelled with bimane showing shifted response to higher proton concentration in comparison with GLIC-Bim136-Q101W. pH applications are shown above each trace and the horizontal scale represent 1 minute of recording. Graphs represent pH-dependent curves showing a shift to higher proton concentrations in electrophysiological responses (C) and fluorescence quenching responses (D) for both mutants.

Another mutation, Y28F two residues apart, was reported to produce a moderate gain of function (Nemecz et al., 2017). Surprisingly, mutating Y28F in the Bim136-Q101W-C27S background yields a drastic loss of function characterized by a slow activating receptor and a marked decrease in pH_50_ (Figure 5B and 5C). We verified that this loss of function is due to the combination of C27S and Y28F mutations since, without the endogenous cysteine mutation to serine, Bim136-Q101W-Y28F electrophysiological response is similar that of Bim136-Q101W (Table 1). In fluorescence, the quenching curve of Bim136-Q101W-Y28F-C27S also shows a large decrease in pH_50_, associated with an apparent higher cooperativity. Again, in this case, the ΔpH_50_ are in the same range in fluorescence quenching (−2.2) and in electrophysiology (higher than – 1.5, the plateau could not be reached with this mutant preventing accurate measurement of the pH_50_).

We thus identify here the lower part of the β-sandwich as a region of the protein that controls the quaternary structure of the ECD at pH7 and following pH decrease. The quaternary compaction of the top of the ECD, monitored with the Bim136-Q101W sensor, thus appears strongly coupled in an allosteric manner with the lower part of the ECD interface.

### Long-range allosteric coupling between the TMD and the top of the ECD

To investigate whether allosteric coupling occurs with more distant regions of the protein, we selected three loss of function mutations further away from the Bim136-Q101W pair: D32E nearby the ECD-TMD interface ; H235Q in the middle of the TMD and E222Q, at the bottom of the TMD lining the pore (Figure 6A) (Sauguet et al., 2014) (Nemecz et al., 2017).

**Figure 6.**
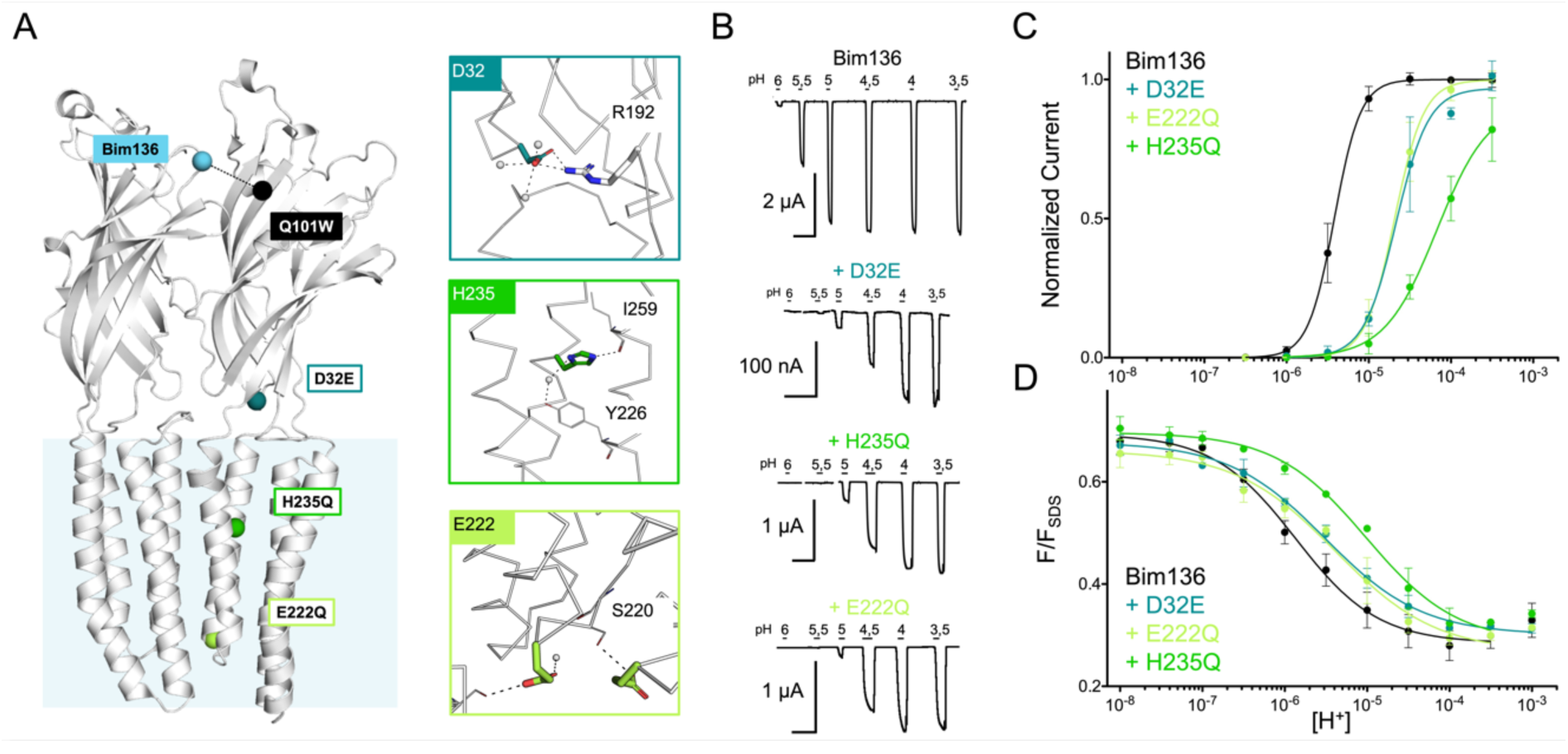
Allosteric coupling between the top of the ECD and around the TMD. (A) Structure of two monomers of GLIC pH 4 showing positions of the fluorescence sensor (Bim136-Q101W) at the top of the ECD and three mutations distributed along the protein. Right panels show zooms on important interactions with the mutated residues. (B) Electrophysiological recordings of the 3 mutants in oocytes, labelled with bimane. Recording of GLIC Bim136-Q101W is shown for comparison. pH applications are shown above each trace and the horizontal scale represent 1 minute of recording. pH-dependent curves for electrophysiological response (C) and fluorescence quenching (D) for the three mutants in comparison with Bim136-Q101W showing a shift to higher proton concentration of the response for all three mutants.

Performing these mutations on the Bim136-Q101W-C27S background shows overall a conservation of their phenotype, with a 10-fold (D32E and E222Q) and more than 30-fold (H235Q) decrease in the pH_50_ of activation as compared to Bim136-Q101W-C27S (Figure 6B and 6C). The fluorescence quenching curves are also shifted to lower pH_50_s, with ΔpH_50_s of 3-fold (D32E and E222Q) and 10-fold (H235Q) (Figure 6D).

The quenching data thus reveal an allosteric coupling between both ends of the protein, since the structural perturbations performed around the TMD are transmitted to the top of the ECD, impairing its compaction. However, as opposed to the ECD mutations E26Q and Y28F/C27S, these mutations have a stronger effect on the pH_50_ of the electrophysiological response as compared to fluorescence quenching. It thus suggests that both processes are not fully coupled for mutations further away from the sensor site.

### Total loss of function mutations decouple ECD and TMD allosteric motions

To further explore the conformational landscape accessible to GLIC, we extended the analysis to mutations known to totally prohibit channel opening (Figure 7A). We selected three mutants: H235F, L157A and L246A which show robust surface expression and very little to no current in oocytes (Figure 7B & 7-Supplementary 1). As electrophysiology could not be used to follow pore opening, we monitored the motion of the M2-M3 loop with Bim250-Y197, in addition to the ECD quaternary and tertiary sensors Bim136-Q10W and Bim135-W72 (Figure 7C).

**Figure 7.**
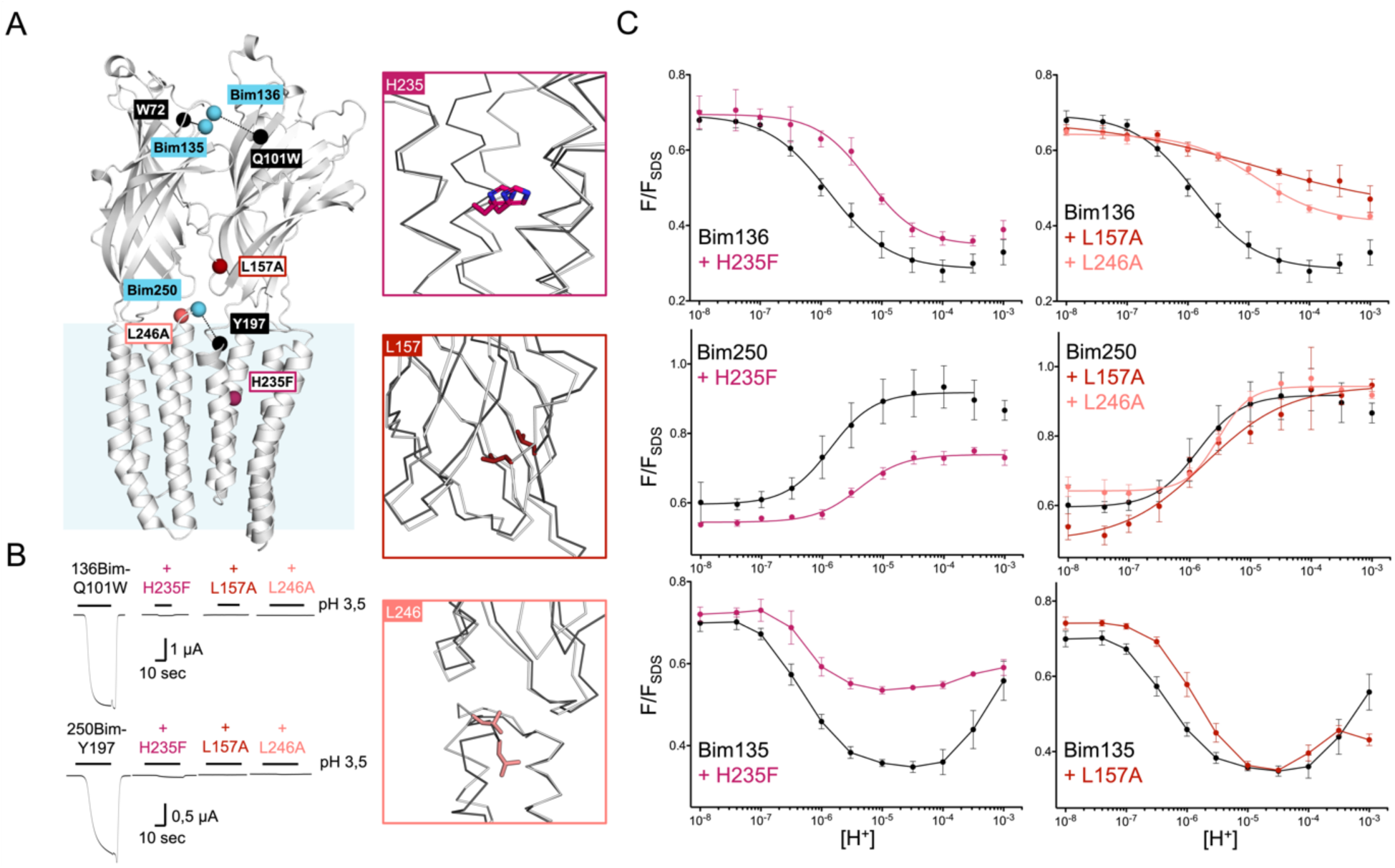
non-functional mutants decouple ECD and TMD motions. (A) Structure of two monomers of GLIC pH 4 showing the position of the fluorescence sensors (Bim136-Q101W, Bim135-W72 and Bim250-Y197) and three mutations causing a total loss of function. Right panels show zooms on important reorganisations of the mutated residues between structures at pH 4 (grey) and pH7 (black). (B) Electrophysiological recordings in oocytes of the 3 mutants labelled with bimane showing no current in comparison with GLIC presenting sensor mutations only. (C) pH-dependent curves in fluorescence for the three mutants with the sensor Bim136-Q101W, Bim250-Y197 and Bim135-W72.

Mutants L157A and L246A reveal unique quenching phenotypes. Combined with Bim136-Q101W, they both show a pH-dependent quenching of fluorescence with a greatly decreased amplitude associated with a significant decrease in pH_50_ as compared to the Bim136-Q101W background. When combined with Bim135-W72, L157A shows a decrease in pH_50_ of the first quenching component, and a diminished amplitude of the second unquenching component. In contrast, neither L157A nor L246A significantly alter the movement at Bim250, which occurs with a complete amplitude and no change in pH_50_. Thus, unlike the moderate loss of function mutants investigated above, these mutations decouple the protein motions, partially impairing quaternary compaction of the ECD but allowing full motion of the M2-M3 loop.

The mutation H235F leads to a phenotype opposite to that of L157A or L246A. It allows for a full compaction of the ECD, since its Bim136-Q101W pH-dependent curve shows a full quenching amplitude and a decrease in pH_50_. Combined with Bim135-W72, H235F displays a phenotype with a notably much smaller amplitude of the first component. In contrast, it impairs the movement of the M2-M3 loop, with only a partial pH-dependent dequenching at Bim250. H235F thus seems to decouple the protein motion by partially impairing the M2-M3 motion, but allowing full movement of the ECD.

Altogether, mutants H235F, L157A and L246A lead to quenching phenotypes that do not match any X-ray structures of GLIC WT, suggesting these mutants of GLIC visit new global conformations.

### Allosteric modulator propofol displaces allosteric equilibriums of activation

We also used the TrIQ technique to dissect the mechanism of action of the general anesthetic propofol, an allosteric modulator of GLIC. Propofol binds at three main sites within the TMD: one site in the pore itself nearby the middle of the TMD, and two sites in the upper part of the TMD at intra-or inter-subunit locations (Figure 8A). Propofol is an inhibitor of GLIC, but it has been shown to be a potentiator of the H235Q mutant (Fourati et al., 2018). We verified that these effects are conserved in the Bim136-Q101W-C27S background, with propofol decreasing the pH_50_ of activation of Bim136-Q101W while increasing the pH_50_ of Bim136-Q101W-H235Q (Figure 8B and C). Interestingly, on both mutants, fluorescence quenching experiments essentially parallel the electrophysiological data. Addition of 100 μM propofol on Bim136-Q101W decreases the fluorescence pH_50_ by half a unit, while it increases that of Bim136-Q101W-H235Q by more than half a unit (Figure 8D). Data thus show that propofol do not act locally by altering the conformation of the TMD, but rather acts on the global allosteric transitions by displacing the equilibrium between resting and active conformation and preserving coupling between ECD and TMD.

**Figure 8.**
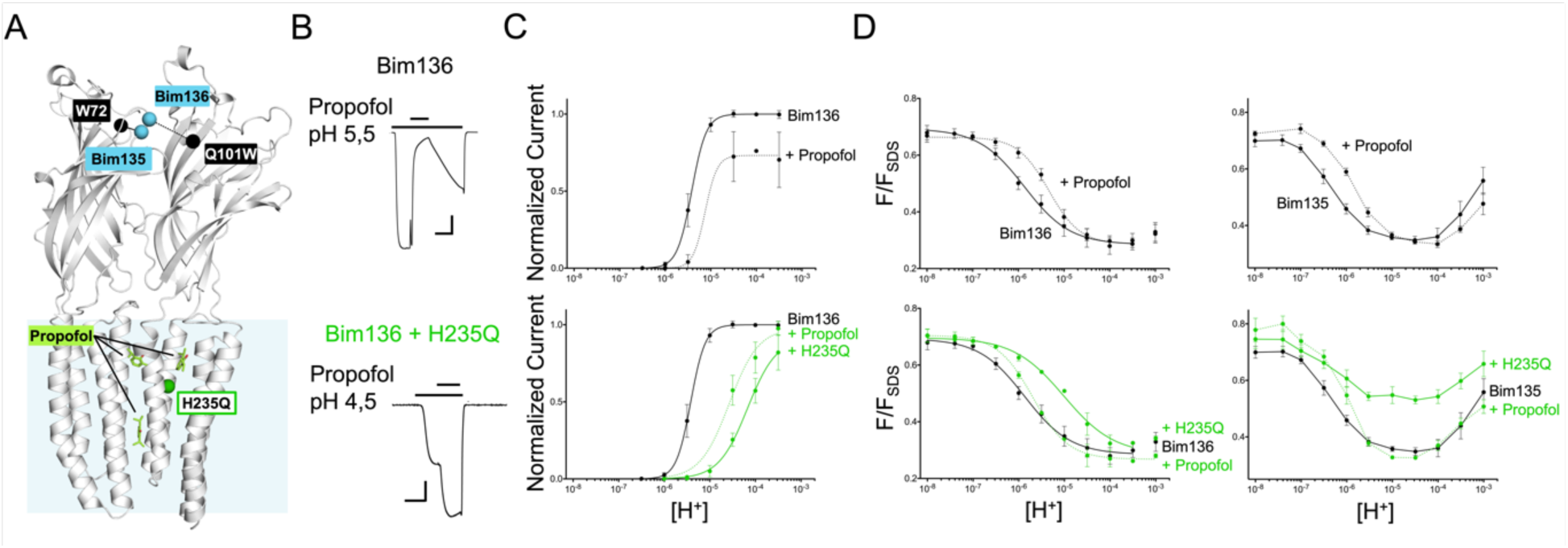
Effect of propofol on ECD compaction. (A) Structure of two monomers of GLIC pH 4 showing positions of the fluorescence sensor Bim136-Q101W at the top of the ECD and three propofol binding sites intra, inter-subunit and in the pore identified by X-ray crystallography. Example of electrophysiological response to 100 μM propofol during a low pH application (B) and dose response curves (C) in Bim136-Q101W with and without the H235Q mutation showing inhibition and potentiation respectively (scale bars represent 100 nA and 30 sec). (D) Effect of 100 μM propofol on fluorescence quenching without (top panels) and with H235Q mutation (lower panels) for the Bim136-Q101W and Bim135-W72 sensors.

We also investigated propofol in the Bim135-W72 H235Q mutant. Interestingly this mutant shows a similar quenching curve to the one of Bim135-W72 H235F, with the difference that electrophysiological response can be measured with mutation in Gln while it is not functional in Phe. The fluorescence recordings show that propofol elicits a decrease in pH_50_ for the first quenching component of Bim135-W72. For Bim135-W72-H235Q, the quenching curve displays a much lower amplitude as compared to the WT, while propofol restores a WT-like pH-dependent quenching curve. The observed global lower amplitude in Bim135-72 H235Q/F mutant may either result from a decrease in the reorganization responsible for the amplitude of first component, or a rightward shift of the curve resulting in an overlap and therefore averaging of the bimane quenching and unquenching curves. Therefore, the fluorescence data related to the Bim135-W72 pair cannot be interpreted in structural terms, however they further document the strong allosteric coupling with the upper part of the ECD.

## Discussion

### Long-range allosteric coupling associated with pre-activation and pore-opening processes

In this study, the fluorescence quenching and electrophysiological data are recorded on an ensemble of GLIC proteins, on which pH-dependent changes lead to a cascade of allosteric reorganizations from the resting to the active and potentially desensitized conformations.

We found that a series of five loss-of-function mutations, which shift the pH-dependent electrophysiological curves to higher concentrations, also shift the pH-dependent fluorescence quenching curve of ECD-compaction at the extracellular top of the protein. The ECD-compaction is thus sensitive to mutations scattered along the protein structure down to the opposite cytoplasmic end, indicating substantial allosteric coupling.

Our previous kinetic analysis showed that the conformational motions followed by fluorescence occur early in the pathway of activation (Figure 4A). Following a rapid pH-drop using a stopped-flow device, these “pre-activation” motions are complete in less than a millisecond, much faster than pore opening that occurs in the 30-150 millisecond range in electrophysiology (Laha et al., 2013). Since pre-activation precedes pore-opening, it is likely that a shift in pre-activation (followed here by fluorescence) will be reflected as a parallel shift in activation (followed by electrophysiology). ECD mutations E26Q and Y28F/C27S present such a phenotype of paralleled variations in electrophysiology and fluorescence quenching pH_50_ which indicates that those mutations mainly impact the pre-activation transition. In contrast, ECD-TMD interface and TMD mutations D32E, E222Q and H235Q lead to a stronger pH_50_ shift in electrophysiology than in fluorescence quenching. This phenotype suggests that these mutations alter not only the pre-activation, but also the downstream pore-opening transitions leading to an additive effect on the function.

In the same line, the quenching data related to the allosteric modulation of propofol suggest its major effect on the pre-activation transition. The emerging picture is thus that pre-activation, that is associated with complete motions at the various quenching pairs, involves a global reorganization of the protein. The low cooperativity of the fluorescence curves potentially suggests that more than one state might be involved in this process. In contrast, the downstream pore-opening process would involve a more local reorganization that would be restricted to the TMD (E222Q, H235Q, pore opening) and the lower part of the ECD (D32E). We speculate that the slowness of this process may arise from the wetting of the pore, particularly its upper region, which is highly hydrophobic and for which hydration should be energetically costly.

To investigate the structural dynamics of the protein, we generated two *in silico* conformational trajectories between the resting-like and active-like X-ray structures of GLIC. Both trajectories show significant coupling between the opening of the gate in the pore, the motion of the M2-M3 loop, and the contraction of the lower part of the β-sandwich. These structural couplings were already highlighted in a recent all-atom molecular dynamic simulation with a swarm-based string method to solve for the minimum free-energy gating pathways of GLIC activation (Lev et al., 2017). Remarkably, the two trajectories show an inverse sequence of events with a gate opening in the last vs first frames of the transition for trajectories A and B respectively. The pre-activation model described above involves a late gate opening and overall fits trajectory A.

### Conformational flexibility of GLIC during the pre-activation transition

An important finding of this work is that the quenching pattern of the total loss of function mutations revealed novel non-conductive conformations, where the movement of the ECD and TMD are partially or fully decoupled. In GLIC WT, His235 interacts with M3 through a hydrogen bond in the open state (Rienzo et al., 2014). Unlike a mutation to glutamine, mutation of His235 to phenylalanine prevents this critical interaction leading to a non-functional receptor (Prevost et al., 2012). We show here that following a pH decrease H235F undergoes a complete “quenching motion” of the ECD, but only a partial “unquenching motion” of the M2-M3 loop. The mutation thus seems to block the protein in an intermediate state in the A trajectory, where the pH-elicited motions of the ECD are complete but not transmitted to the TMD (Figure 9). Interestingly, the structure of the H235F mutant was previously solved by crystallography at pH 4 in a “locally closed” (LC, 3TLT) conformation (Prevost et al., 2012). This conformation is characterized by a fully active-like structure of the ECD, but a resting-like structure of the TMD including the M2-M3 loop and a closed channel. This structure fits well with the quenching data of H235F at Bim136-Q101W, but not with that of Bim250-Y197 where the fluorescence quenching data indicate a partial movement of M2-M3 that is not seen in the X-ray structure. Inspection of the structure of H235F shows however a movement of the quencher Y197 which side chain is re-oriented toward the transmembrane domain, away from Bim250, a feature that could plausibly account for this partial unquenching.

**Figure 9.**
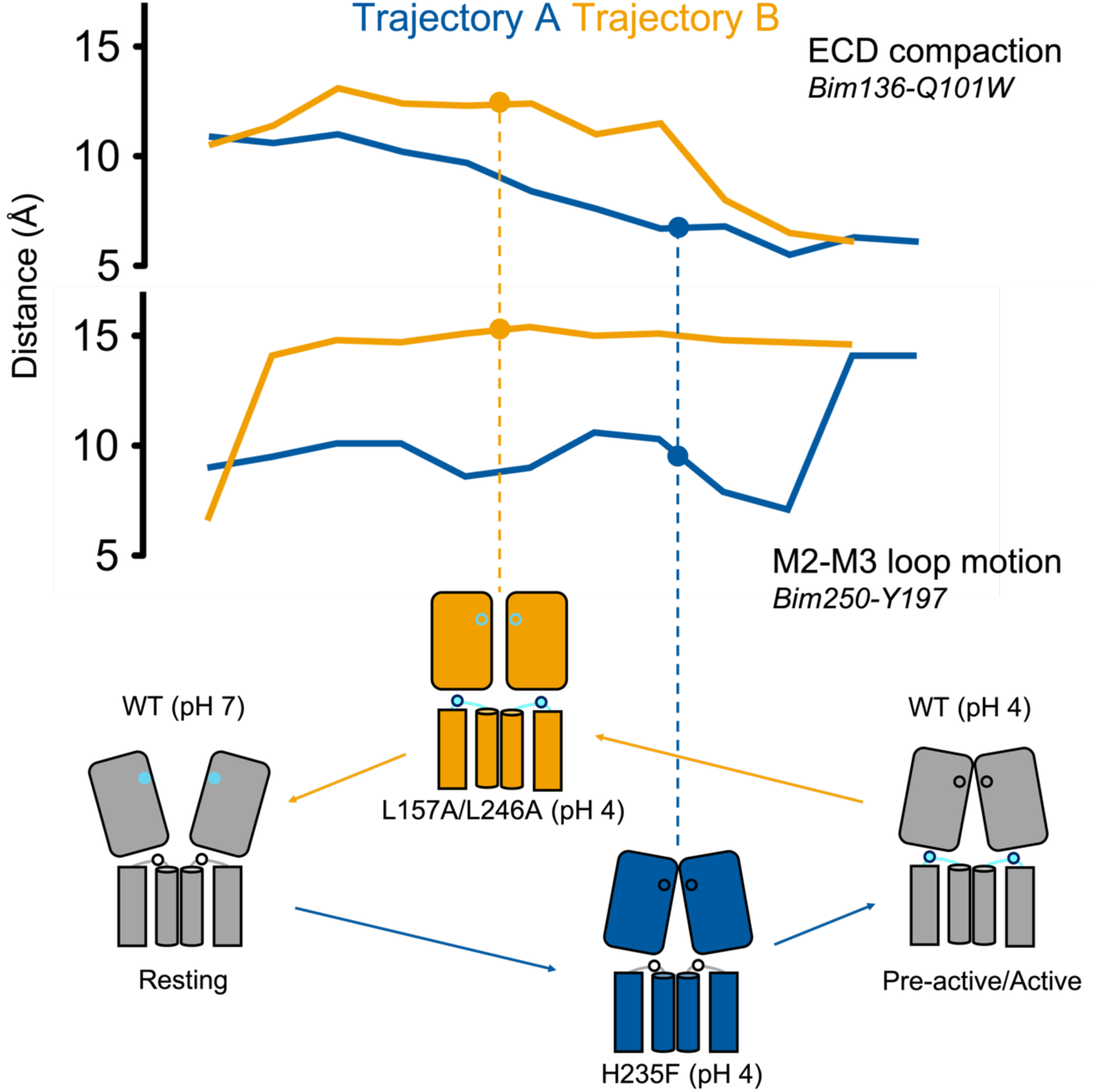
Speculative summary scheme. Bimane-quencher centroid distances are shown for the trajectories A and B. Non-functional mutants H235F and L157A/L246A are tentatively placed on the trajectories A and B respectively, according to their quenching pattern.

Leu157 and Leu246, located around the ECD-TMD interface, are involved in strong hydrophobic interactions in the active state, Leu157 stabilizes a compact conformation of the β-sandwich, and Leu246 a key ECD-TMD interaction (Sauguet et al., 2014). Mutations L157A and L246A display an inverse phenotype as compared to H235F, since they allow a complete “unquenching” motion of the M2-M3 loop but a partial compaction of the ECD. These mutations thus seem to block the protein in an intermediate state within the trajectory B, and this time, with the complete pH-elicited motions of the M2-M3 loop, but not completely transmitted to the top of the ECD nor the channel gate. Altogether, these mutations reveal key regions that act as hinges within the signal transduction pathways of GLIC (Mowrey et al., 2013).

It is noteworthy that the mechanism whereby protons activate GLIC remains elusive. In particular, systematic mutational analysis of every residues that change protonation state in the pH4-6 range (Asp, Glu and His) indicate that protons act on several sites to activate GLIC, with a particularly strong effect of mutations nearby the ECD-TMD interface (Nemecz et al., 2017; Hu, Ataka, et al., 2018). With the three mutations H235F, L157A and L246A, we show here that pH change elicits motions independently at two levels, the ECD compaction (in H235F) and the M2-M3 loop movement (in L157A and L246A), further supporting such a multi-sites effect.

Another important conclusion from our data is that the protein has access to an unanticipated repertoire of conformations, and NMA analysis suggests that these conformations might contribute to different allosteric pathways of pre-activation. The idea that GLIC can follow different trajectories during the gating process was already proposed. For instance, through the use of a hybrid elastic-network Brownian dynamics simulation predicting two possible pathways for GLIC gating, that are characterized by different compactions of the ECD (Orellana et al., 2016). Here, we extend this concept by proposing two pathways involving either an early motion of the ECD or an early motion of the M2-M3 loop (Figure 9).

### Consequences on the gating mechanism within the pLGIC family

The conservation of the gating mechanism between bacterial and eukaryotic pLGICs is well documented by the available structures with the common allosteric regulatory sites for ligands and mutations (Sauguet et al., 2015; Bertozzi et al., 2016; Rienzo et al., 2016), together with the allosteric compatibility between eukaryotic and prokaryotic ECD/TMD domains to form functional chimeras (Duret et al., 2011; Moraga-Cid et al., 2015; Laverty et al., 2017). It is therefore tempting to speculate that the pre-activation transition of GLIC that we characterize here might have counterparts in human neurotransmitter-gated receptors. In this line, some recent structures of eukaryotic receptors including the 5-HT_3_R (Polovinkin et al., 2018) and the GABA_A_R (Masiulis et al., 2019) show pre-active-like conformations characterized by marked agonist-elicited reorganization of the ECD but a closed channel at the TMD. Additionally, the flipped state, where the conformational change of the orthosteric site is predicted to be rather complete, but where the channel is closed, would fit the functional requirement of a pre-active state (Lape et al., 2008). Our observation that propofol specifically affects the pre-activation step might thus tentatively be extended to eukaryotic receptors.

An unexpected finding here is that GLIC and its mutants have access to a large repertoire of conformational states. It raises the possibility that GLIC can follow several conformational trajectories during the gating transitions. This idea challenges the conventional concept that receptor activation involves a single conformational pathway. For instance, REFER analysis of the muscle-type nAChR show that two discrete regions undergo an early motion during activation, the orthosteric site and the M2-M3 loop. These data were interpreted in the framework of a four-state linear transition pathway (Gupta et al., 2017), but alternatively, it is plausible that the protein can follow two pathways, one starting at the orthosteric site and the other at the M2-M3 loop.

Our work also investigates the mechanism of action of allosteric mutations by measuring their effects at different levels of the protein. The loss of function mutations that shift the pH-dependent activation curves produce parallel shifts in the fluorescence quenching, indicating that they alter the equilibrium constant “L” between the allosteric states involved in activation, driving them toward the resting state (Galzi et al., 1996). In sharp contrast, the total loss-of-function mutants, silent in electrophysiological recordings, were found to undergo proton-elicited allosteric transitions by fluorescence quenching, visiting other conformations that plausibly correspond to intermediates blocked within the activation pathway. Actually, allosteric mutations of neurotransmitter-gated receptors, causing congenital pathologies including myasthenia and hyperekplexia have been extensively studied (Taly and Changeux, 2008; Bode and Lynch, 2014; Hernandez and Macdonald, 2019). Most of the hot spots mutated here on GLIC were found associated with pathologies on human receptors. In particular, the lower part of the ECD-ECD interface is the site of a *de novo* S76R mutation in GABA_A_ α1 (homologous to Glu26) causing epilepsy (Johannesen et al., 2016) and the mutation L42P in the nAchR d (homologous to Cys27) causing myasthenia (Shen et al., 2008). This latter mutation (as well as mutation of Asn41, homologous to Glu26) decreases activation kinetics and this residue was shown to be energetically coupled to Tyr127 on the other side of the interface. Interestingly, equivalent residues in GLIC (Cys27; Glu26 and Tyr111) are part of a water network at the bottom of the ECD (Figure 5A). Another notable example is the mutation P250T in GlyR α1 that causes hyperekplexia (Saul et al., 1999) and which is homologous to Glu222 in GLIC. Our work on GLIC gives general mechanisms of how mutations affect pLGICs transitions, and further work, for example by voltage-clamp fluorometry, would be required to challenge such mechanisms in the context of congenital pathologies on neurotransmitter receptors.

## Material and methods

### Mutagenesis

All GLIC mutants were obtained using site directed mutagenesis on the C27S background of GLIC, except for the Bim136-Y28F-C27 where the endogenous cysteine was introduced back. Similarly, to previous studies, two different vectors were used: a pet20b vector with GLIC fused to MBP by a linker containing a thrombin cleavage site under a T7 promoter for expression in E. coli BL21; a pmt3 vector for expression in oocytes with GLIC containing a Cter HA tag and in Nter the peptide signal from alpha7-nAChR. Incorporation of the mutations in both vectors were verified by sequencing.

### GLIC mutant production and purification

Protein production of MBP-GLIC and labeling was done as previously described (Menny et al., 2017) with a few modifications. In brief, MBP-GLIC was expressed in BL21 *E coli* cells overnight at 20°C after induction by 100 µM IPTG. Cells were collected and resuspended in buffer A containing 20 mM Tris; 300 mM NaCl at pH 7.4 and subsequently disrupted by sonication. After membrane separation by ultracentrifugation, membrane proteins were extracted overnight in buffer A supplemented with 2 % DDM. After ultracentrifugation, supernatant was incubated with amylose resin and MBP-GLIC was eluted using buffer A supplemented with DDM 0.02 % and saturating concentration of maltose. To remove endogenous maltoporin contaminant, a first size exclusion chromatography was performed on superose 6 10/300 GL in buffer A with 0.02 % DDM. GLIC-MBP concentration was measured and the protein was incubated overnight at 4°C with thrombin to cleave off MBP and with monobromobimane (mBBr) at a 1:5 (GLIC monomer:fluorophore) ratio, to label the protein. The mBBr dye being solubilized in DMSO, the sample volume was adjusted to remain below 1% DMSO final concentration. After labeling, a second gel filtration was done to get rid of the MBP and unbound dye molecules. GLIC-Bimane samples were flash frozen in liquid nitrogen and stored at -80°C prior to fluorescence measurements.

### Steady-state fluorescence measurements

Fluorescence measurements were done as previously described (Menny et al., 2017). Samples were equilibrated to room temperature and diluted with buffer A with 0.02 % DDM to reach a concentration around 40 µg.mL^-1^. Fluorescence recording buffers consisting of 300 mM NaCl, 2.7 mM KCl, 5.3 mM Na_2_HPO_4_ and 1.5 mM KH_2_PO_4_ were prepared beforehand and their pH was adjusted either to 7.4 or to different pH in order to reach the desired pH value (from pH 8 to 3) after mixing equal volumes with Buffer A 0.02 % DDM. Measurements were done at 20°C in 1 mL disposable UV transparent 2.5 mL cuvettes in a Jasco 8200 fluorimeter with 385 nm excitation wavelength and the emission spectra was recorded through 2.5 nm slits from 420 to 530 nm. Other parameters were kept constant throughout the study. On the sample at pH 7.4, an addition of SDS to reach 1 % final concentration was done to obtain the F_SDS_ value and a tryptophan emission spectrum was done before and after SDS addition in order to monitor denaturation.

Fitting of fluorescence measurements was done on each fluorescence series (values from one pH range) with at least 3 series per mutant using the following Hill equation:

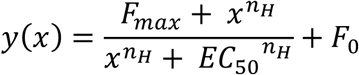

where F_max_ represents the maximal change in fluorescence amplitude; F_0_ the initial fluorescence at pH 7.8; n_H_ represents the hill number and EC_50_ the proton concentration for which half of the maximal fluorescence change is measured. For Bim136-Q101W and Bim250-Y197 and in some other mutants, we excluded the data point below pH 3.5 that show a small but significant change in fluorescence intensity in the opposite direction to the quenching curves. We did not fit the Bim135-W72 mutant that shows a bell shape curve.

### Electrophysiological recordings

Electrophysiological recordings of GLIC were made on *Xenopus* oocytes provided either by the Centre de Ressources Biologiques Xénopes (Rennes-France) or by Ecocyte Bioscience (Dortmund-Germany). Recordings were made as previously described (Nury et al., 2011) with oocytes 48-96 h post nucleus injection with a mix containing 80 ng.µL^-1^ of GLIC cDNA and 25 ng.µL^-1^ of GFP cDNA. Recording were done in MES buffer containing 100 mM NaCl, 3 mM KCl, 1 mM CaCl2, 1 mM MgCl2 and 10 mM MES with pH adjusted by addition of 2 M HCl. The perfusion chamber contained two compartments and only a portion of the oocyte was perfused with low pH solution. Bunte salt bimane labeling was performed prior to recording by incubation for 1h at room temperature with the dye concentrated at 1 mM in MES buffer. To correct data for rundown, a solution with a pH value in the middle of the dose response (usually pH 5) was used as a reference at the beginning and the end of the dose response and every 3/4 applications. To limit the effect of propofol that can stay in the membrane in-between applications (Heusser et al., 2018), only a limited number of pH solution were tested per dose-response.

Electrophysiological recordings were analyzed using AxoGraph X and Prism was used to fit individual dose responses using the Hill equation:

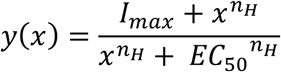

where I_max_ represents the maximal current in percentage of the response from the reference solution. n_H_ represents the hill number and EC_50_ the proton concentration for which half of the maximal electrophysiological response is recorded.

### Xenopus oocytes immunolabeling

Mutants generating currents smaller than 500 nA at high proton concentrations were categorized as non-functional. For these non-functional mutants, expression tests were performed by immunolabeling of oocytes as previously described (Prevost et al., 2012; Sauguet et al., 2014). 3 to 4 days post injection, GFP positive oocytes were fixed overnight in paraformaldehyde (PFA) 4 % at 4°C. Immunolabeling was performed after 30 min saturation by 10 % horse serum in PBS buffer. Rabbit anti HA-tag primary antibody was incubated for 90 min in 2 % horse serum and the secondary antibody anti-Rabbit coupled to Alexa Fluor 645 was incubated for 30 min. After a second PFA fixation overnight, oocytes were included in warm 3 % low-melting agarose and 40 µm slices were made using a vibratome on a portion of the oocyte. Several slices per oocytes were mounted one a slide and analyzed in an epi-fluorescence microscope using constant exposure time between none functional mutant and functional mutants used as positive controls.

### Molecular Modeling

Each structure (4NPQ and 4HFI) was fitted, using iMODfit (Lopéz-Blanco and Chacón, 2013), to the simulated electron-microscopy envelope of the other structure. The EM density map resolution was set to 5 Å and the grid size to 0.5 Å. iMODfit was then used to fit each structure to the density of the other structure with half of the modes considered for the conformational change, yielding trajectory A (4NPQ to 4HFI) and trajectory B (4HFI to 4NPQ). The structure of the protein and ligand were converted to pdbqt files with the software open babel 2.4.1. Covalent docking was then performed with the software smina (Koes et al., 2013). The box for docking has been defined around the mutated residue, with a size of 30 Å in each direction. Covalent docking forced the bimane to be in direct contact with the SG atom of the cysteine which was introduced experimentally.

## Acknowledgments

The work was supported by the ‘Agence Nationale de la Recherche’ (grant ANR-13-BSV8-0020, Pentagate), the doctoral school ED3C and the ‘Foundation pour la Recherche Médicale’ (PhD funding to SNL), the “Initiative d’Excellence” (cluster of excellence LABEX Dynamo, ANR-11-LABX-0011 to AT) and the ERC (grant No. 788974, Dynacotine). The authors would like to thank Marc Gielen, Akos Nemecz and Marie Prévost for critical reading of the manuscript.

## Author contribution

SNL and PJC designed fluorescent quenching experiments; SNL, AM and KM performed experiments; AT designed and performed in silico simulations. All authors analyzed the data. SNL and PJC wrote the manuscript with the help of the other authors.

## Supplementary figures

**Figure 3-Supplementary 1.**
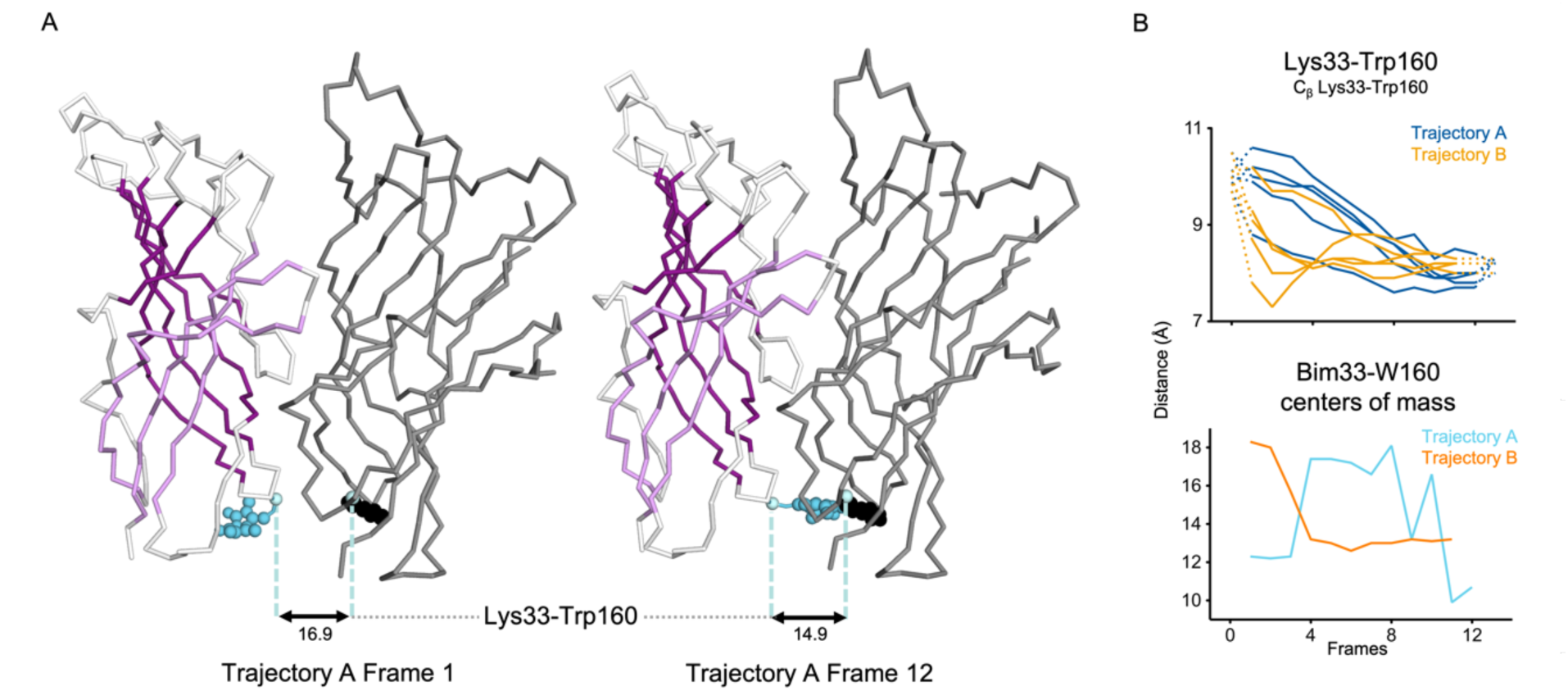
Evolution of ECD inter-subunits distances at the bottom of the ECD in iMODfit trajectories. (A) Snapshots of two subunits of GLIC ECD in the first and last frame of the trajectory A with Bim33-W160 quenching pair modeled. One subunit is shown in grey, the other in white with β-sheets forming the β-sandwich shown in dark and light purple; bimane is shown in blue and quencher in black spheres; C_β_ atoms used for measurements are shown in pale blue spheres and distances are indicated in angstroms. (B) Inter-subunit distances showing ECD compaction at the Lys33-Trp160 level (top panel) and between bimane and W160 centers of mass (bottom panel) in both trajectories A and B. Points at frames 0 and 13 are the distances in pH4 and pH7 structures subunits interfaces.

**Figure 3-Supplementary 2.**
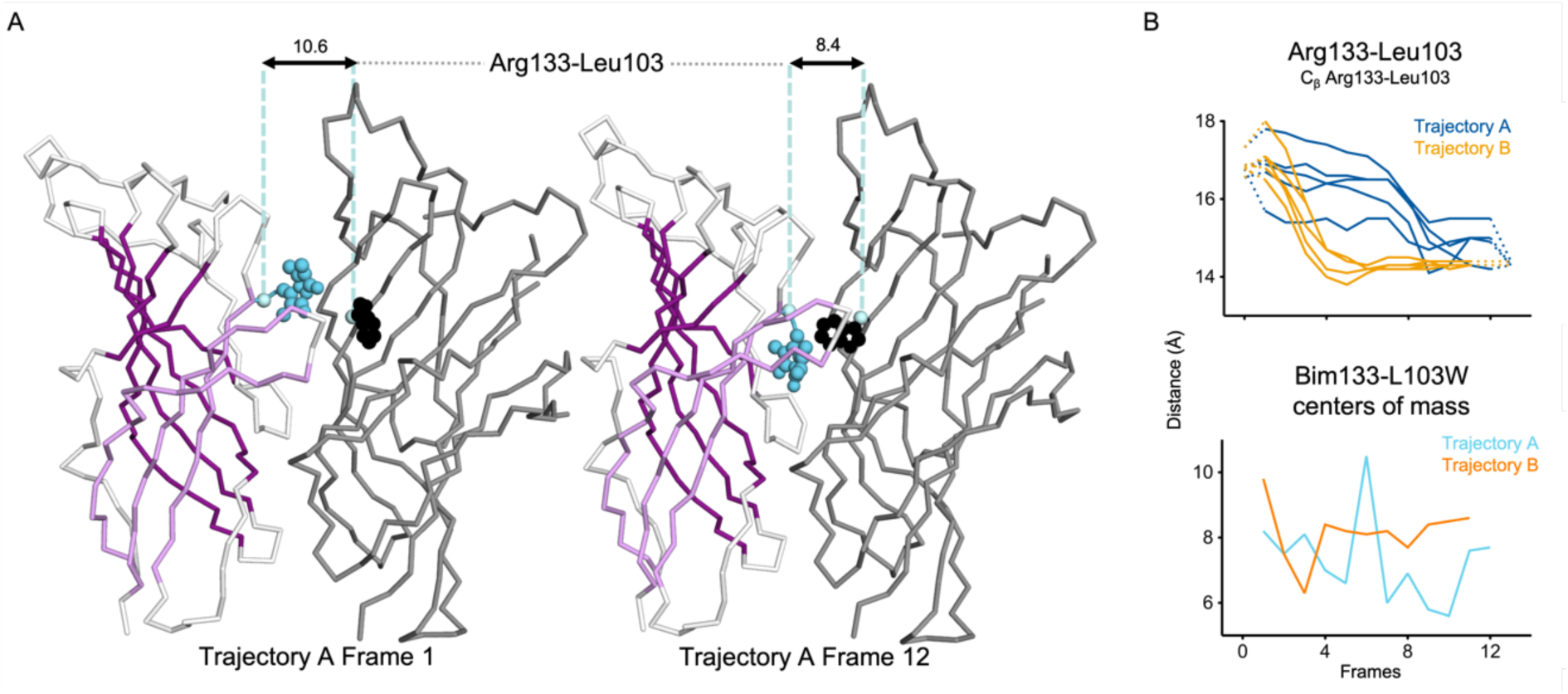
Evolution of ECD inter-subunits distances at the top of the ECD in iMODfit trajectories. (A) Snapshots of two subunits of GLIC ECD in the first and last frame of the trajectory A with a Bim133-L103W quenching pair modeled. One subunit is shown in grey, the other in white with β-sheets forming the β-sandwich shown in dark and light purple; bimane is shown in blue and quencher in black spheres; C_β_ atoms used for measurements are shown in pale blue spheres and distances are indicated in angstroms. (B) Inter-subunit distances showing ECD compaction at the Arg133-Leu103 level (top panel) and between bimane and L103W centers of mass (bottom panel) in both trajectories A and B. Points at frames 0 and 13 are the distances in pH4 and pH7 structures subunits interfaces.

**Figure 3-Supplementary 3.**
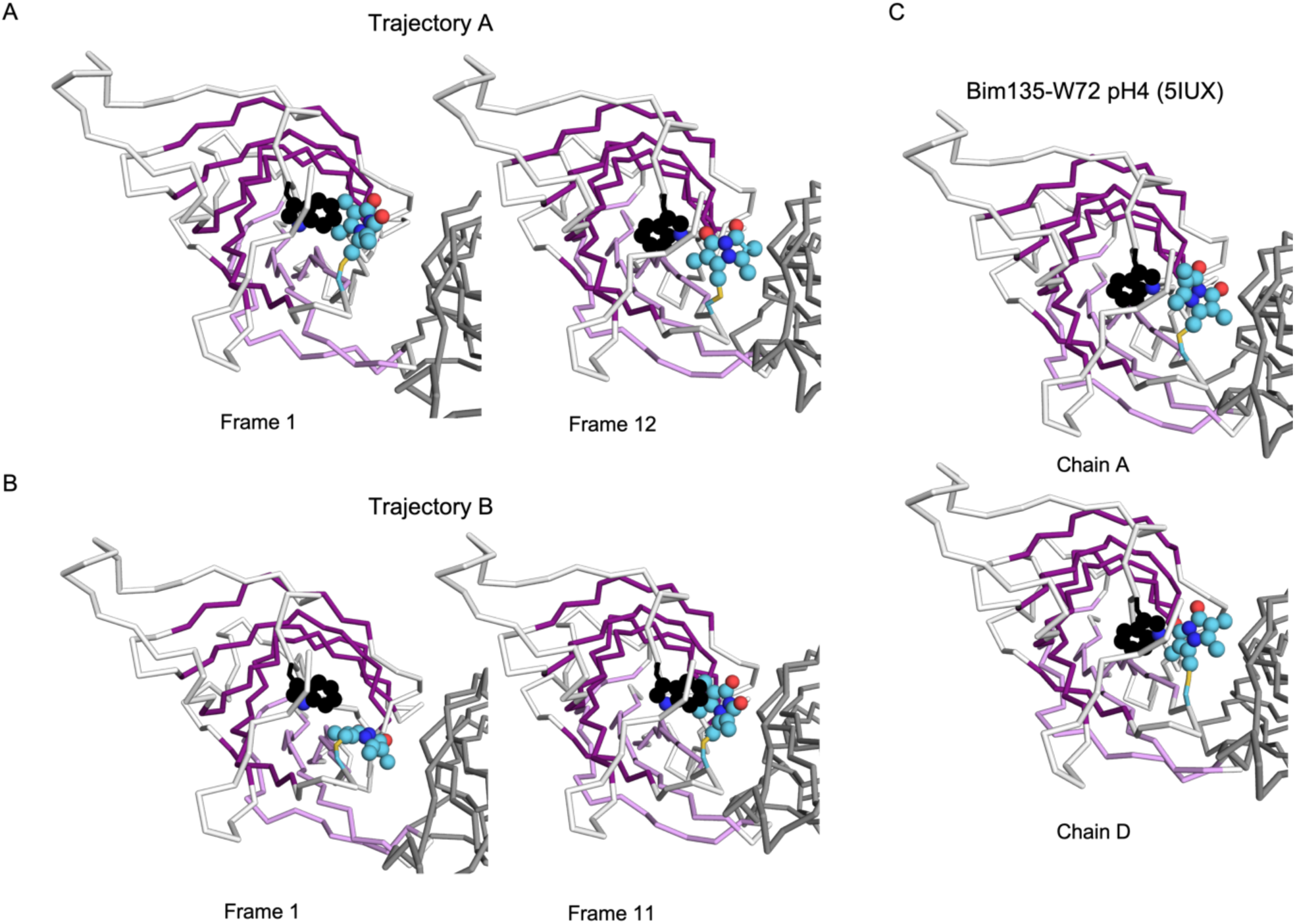
Evolution of Bim135-W72 orientation at the ECD intra-subunits in iMODfit trajectories. (A) Snapshots of one subunit of GLIC ECD, top view in the first and last frame of the trajectory A with Bim135-W72 quenching pair modeled. The adjacent subunit is shown in grey, and the main subunit is shown in white with β-sheets forming the β-sandwich shown in dark and light purple; bimane is shown in blue and quencher in black spheres. (B) Snapshots for the trajectory B. (C) Snapshots from the structure resolved by X-ray of Bim135-W72 at pH 4 (5IUX). Bimane was resolved in two out of five chains and show a similar orientation to the one found in the last frames of both trajectories (A and B right panels).

**Figure 7-Supplementary 1.**
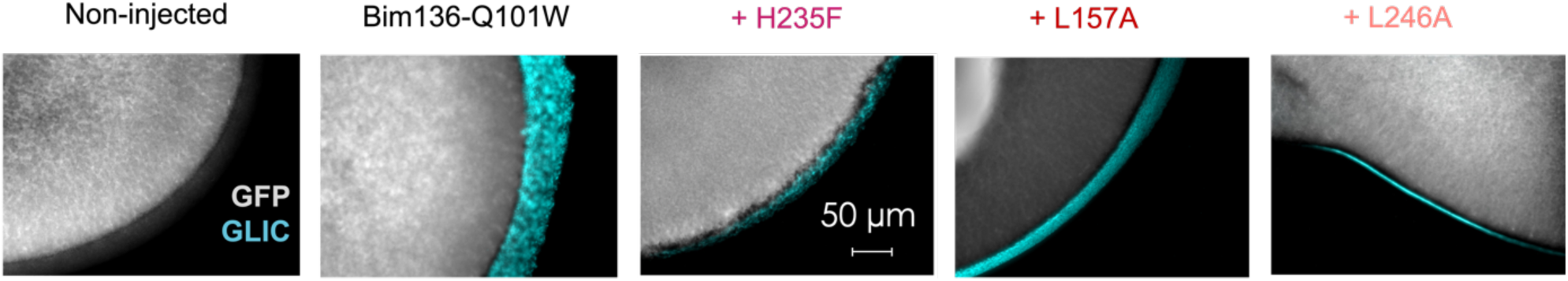
Immunofluorescence microscopy data showing GLIC expression at the oocytes surface. In grey is the GFP fluorescence and in blue the fluorescence resulting from GLIC immunolabeling via anti-HA antibody.

## Notes

### Competing Interest Statement

The authors have declared no competing interest.

